# Consensus interpretation of the Met34Thr and Val37Ile variants in *GJB2* by the ClinGen Hearing Loss Expert Panel

**DOI:** 10.1101/493130

**Authors:** Jun Shen, Andrea M. Oza, Ignacio del Castillo, Hatice Duzkale, Tatsuo Matsunaga, Arti Pandya, Hyunseok P. Kang, Rebecca Mar-Heyming, Saurav Guha, Krista Moyer, Christine Lo, Margaret Kenna, John Alexander, Yan Zhang, Yoel Hirsch, Minjie Luo, Ye Cao, Kwong Wai Choy, Yen-Fu Cheng, Karen B. Avraham, Xinhua Hu, Gema Garrido, Miguel A. Moreno-Pelayo, John Greinwald, Kejian Zhang, Yukun Zeng, Zippora Brownstein, Lina Basel-Vanagaite, Bella Davidov, Moshe Frydman, Tzvi Weiden, Narasimhan Nagan, Alecia Willis, Sarah E. Hemphill, Andrew R. Grant, Rebecca K. Siegert, Marina T. DiStefano, Sami S. Amr, Heidi L. Rehm, Ahmad N. Abou Tayoun, on behalf of the ClinGen Hearing Loss Working Group

## Abstract

**PURPOSE:** Pathogenic variants in *GJB2* are the most common cause of autosomal recessive sensorineural hearing loss. The classification of c.101T>C/p.Met34Thr and c.109G>A/p.Val37Ile in *GJB2* are controversial. Therefore, an expert consensus is required for the interpretation of these two variants.

**METHODS:** The ClinGen Hearing Loss Expert Panel (HL-EP) collected published data and shared unpublished information from participating laboratories regarding the two variants. Functional, computational, allelic, and segregation data were also obtained.

**RESULTS:** The panel reviewed the synthesized information, and classified the Met34Thr and Val37Ile variants according to professional variant interpretation guidelines. We found that Met34Thr and Val37Ile are significantly overrepresented in hearing loss patients, compared to the general population. Met34Thr or Val37Ile homozygotes or compound heterozygotes typically manifest mild to moderate hearing loss. Several other types of evidence also support pathogenic roles for those two variants.

**CONCLUSION:** Resolving controversies in variant classification requires coordinated effort among a panel of international multi-institutional experts to share data, standardize classification rules, review evidence, and reach a consensus. The ClinGen HL-EP concluded that Met34Thr and Val37Ile variants in *GJB2* are pathogenic for autosomal recessive nonsyndromic hearing loss with variable expressivity and age-dependent penetrance.

## INTRODUCTION

Variants with incomplete penetrance for Mendelian conditions pose significant challenges for clinical interpretation because they are relatively common in the population and present in healthy individuals^1^. No such variants have been rigorously reviewed and classified according to the American College of Medical Genetics and Genomics and the Association for Molecular Pathology (ACMG/AMP) guidelines^2^. To demonstrate the best practice to interpret such variants, the ClinGen Hearing Loss Expert Panel (HL-EP) applied the ACMG/AMP guidelines to interpret two controversial variants in *GJB2*.

*GJB2* encodes connexin 26, a member of the gap junction protein family. Gap junctions are intercellular channels allowing the coupling of adjacent cells to share molecules, ions, and electrical signals. Each gap junction channel is composed of two connected hemichannels called connexons, one on either membrane of the neighboring cells. Each connexon is a hexamer of the same or different connexin units. Biallelic pathogenic variants in *GJB2* (NM_004004.5) are the most frequently identified genetic cause of autosomal recessive sensorineural hearing loss^3,4^. Hundreds of *GJB2* variants have been reported in patients with hearing loss. Premature termination codons (PTC) of *GJB2*, such as c.35delG, c.167delT, and c.235delC common in European, Ashkenazi Jewish, and Asian populations respectively, are established pathogenic variants. However, classifications of two notable missense variants c.101T>C/p.Met34Thr and c.109G>A/p.Val37Ile have been controversial. Met34Thr was first reported as being associated with dominant hearing loss^3^, but its pathogenicity for dominant hearing loss was later challenged because of subsequent identification of its occurrence in individuals with normal hearing^4,5^, and an autosomal recessive mode of inheritance was suggested^6-8^. Val37Ile was first identified as a polymorphism in a heterozygous control^9^, and later found in the homozygous state or *in trans* with known pathogenic *GJB2* variants in affected individuals^6,10^. Both variants were found relatively frequently in the general population. Some individuals who are homozygotes for these variants appeared to have normal hearing^11,12^. Reduced penetrance has been proposed to explain the inconsistency^13^. We conducted a survey of clinical laboratories in the United States and Canada regarding the classification of these variants, and found significant variability in the classifications across different laboratories (Supplementary Information). Therefore, we consider it a priority to resolve the controversy and reach a consensus.

We collected data and evaluated the evidence for the classification of these two variants. Herein, we report the ClinGen and ClinVar effort to resolve controversies regarding the Met34Thr and Val37Ile variant classification assertions, and demonstrates the best practice to interpret variants with incomplete penetrance.

## MATERIAL METHODS

The ClinGen Hearing Loss Clinical Domain Working Group established a variant curation expert panel, hereafter referred to as the HL-EP, including otolaryngologists caring for patients with hereditary hearing loss, medical geneticists, clinical laboratory diagnosticians, molecular pathologists, genetic counselors, and investigators specialized in auditory research^14^. Data collected from genetic testing laboratories and clinics included total number of tested hearing loss patients, their ethnicities, genotypes, and phenotypic information (age at onset of hearing loss, age at testing, type of hearing loss, severity, laterality, frequency range affected, family history, and other clinical features) of homozygotes and compound heterozygotes with Met34Thr or Val37Ile. Data were analyzed and interpreted according to the ACMG/AMP standards and guidelines for the interpretation of sequence variants^2^ and the criteria recently specified by the ClinGen HL-EP^14^.

### Statistical analysis

For 2x2 contingency tables, odds ratios (OR), 95% confidence intervals (CI), z statistics, and p values were calculated using MEDCALC (https://www.medcalc.org/calc/odds_ratio.php). For comparisons of the severity among different genotype groups, an ordinal logistic regression model was used. No covariates such as age and sex were included, because the information was not available for all individuals. The proportions of the sample sizes of the genotype groups were used as weights for the analysis. Type 1 test was used to test the overall significant difference among all groups and ad-hoc analysis was performed to compare the estimated odds ratio between each pair of specific gene types. The statistical significance was defined as p value <0.05. The analysis was performed with SAS software (version 9.4).

## RESULTS

### Population Data

Met34Thr and Val37Ile have been reported in patients with hearing loss in various populations (Supplementary Information). To perform accurate case-control comparisons, the HL-EP decided not to rely on published cases in the literature due to concerns of the publication bias. Instead, we obtained data from 15 contributing sites (Table 1), including 14 with diagnostic test data on hearing loss patients and six with population or familial screening test data. The level of details provided by each site varied. Only data from sites which provided the total number of unselected probands with hearing loss were included in the statistical analysis. We also used information from the Genome Aggregation Database (gnomAD; http://gnomad.broadinstitute.org/) to represent the general population. Three sites (BCH, DY, and GDWC) reported only cases with biallelic *GJB2* variants involving Met34Thr or Val37Ile. Eleven sites reported results from 17,635 probands with hearing loss tested for these two variants. Methods included specific variant testing, *GJB2* sequencing, large multigene panels, and exome sequencing. Ethnicity information was available from eight sites on 7,962 European probands and 2,066 Asian probands. Five sites (Counsyl, CUHK, NTMC, TAU, and UNC) provided population screening data from 664,114 individuals who tested for these two variants in *GJB2*, including 306,982 Europeans and 66,423 Asians (Table 2).

**Table 1.** Contributing sites. Symbols, names, geographical locations, tested populations, and the type of tests performed are listed.

| **Symbol** | **Name** | **Country /Region** | **Cohort** | **Type** | **All<sup>1</sup>** | **Ethnicity<sup>2</sup>** | **Biallelic<sup>3</sup>** |
| --- | --- | --- | --- | --- | --- | --- | --- |
| BCH | Boston Children's Hospital | USA | Individuals with hearing loss | Diagnostic | 0 | 0 | 35 |
| CCHMC | Cincinnati Children's Hospital Medical Center | USA | Individuals with hearing loss | Diagnostic | 1491 | 692 | 63 |
| CHOP | Children's Hospital in Philadelphia | USA | Individuals with hearing loss | Diagnostic | 163 | 121 | 13 |
| Counsyl | Counsyl | USA | Reproductive adults | Screening | 654426 | 364788 | 0 |
| CUHK | Chinese University of Hong Kong | Hong Kong | Individuals with hearing loss; general population | Diagnostic and screening | 5839 | 5839 | 6 |
| DY | Dor Yeshorim | USA, Israel | Individuals with hearing loss and family members | Diagnostic and screening | 0 | 0 | 18 |
| EGL | EGL Genetics | USA | Individuals with hearing loss | Diagnostic | 1113 | 0 | 32 |
| GDWC | Guangdong Women and Children Hospital | China | Individuals with hearing loss; reproductive adults | Diagnostic and screening | 0 | 0 | 27 |
| HRyC | Hospital Ramon y Cajal | Spain | Individuals with hearing loss | Diagnostic | 3881 | 3777 | 65 |
| IG | Integrated Genetics, LabCorp | USA | Individuals with hearing loss | Diagnostic | 2883 | 0 | 0 |
| LMM | Laboratory for Molecular Medicine, Partners Personalized Medicine | USA | Individuals with hearing loss | Diagnostic | 3472 | 1697 | 135 (1) |
| NTMC | National Hospital Organization Tokyo Medical Center | Japan | Individuals with hearing loss | Diagnostic and screening | 1884 | 1884 | 56 (2) |
| TAU | Tel Aviv University (including those tested at Rabin and Sheba Medical Centers) | Israel | Individuals with hearing loss; reproductive adults | Diagnostic and screening | 719 | 0 | 3 (2) |
| UNC | University of North Carolina | USA | Individuals with hearing loss; general population | Diagnostic and screening | 5833 | 4590 | 8 |
| VGH | Taipei Veterans General Hospital | Taiwan | Individuals with hearing loss | Diagnostic | 45 | 45 | 11 |
| **Total** |  |  |  |  | **680985** | **383388** | **472 (5)** |

1. The total number of affected probands and screened populations contributed to the statistical analysis
2. The total number of probands with relevant ethnicity information contributed to the statistical analysis
3. The total number of Met34Thr or Val37Ile homozygotes and compound heterozygotes with clinical information contributed to the analysis. The numbers of unaffected individuals confirmed by audiology evaluation are in parentheses.

We compared frequencies of individuals with these two variants between cases (patients with hearing loss in our multicenter cohort) and the general population. Met34Thr and Val37Ile homozygotes are significantly enriched in cases over the general population with ORs of 16 (95%CI 11-25, Z=13, p<0.0001) and 20 (95%CI 17-24, Z=31, p<0.0001) respectively (Table 2).

**Table 2.**
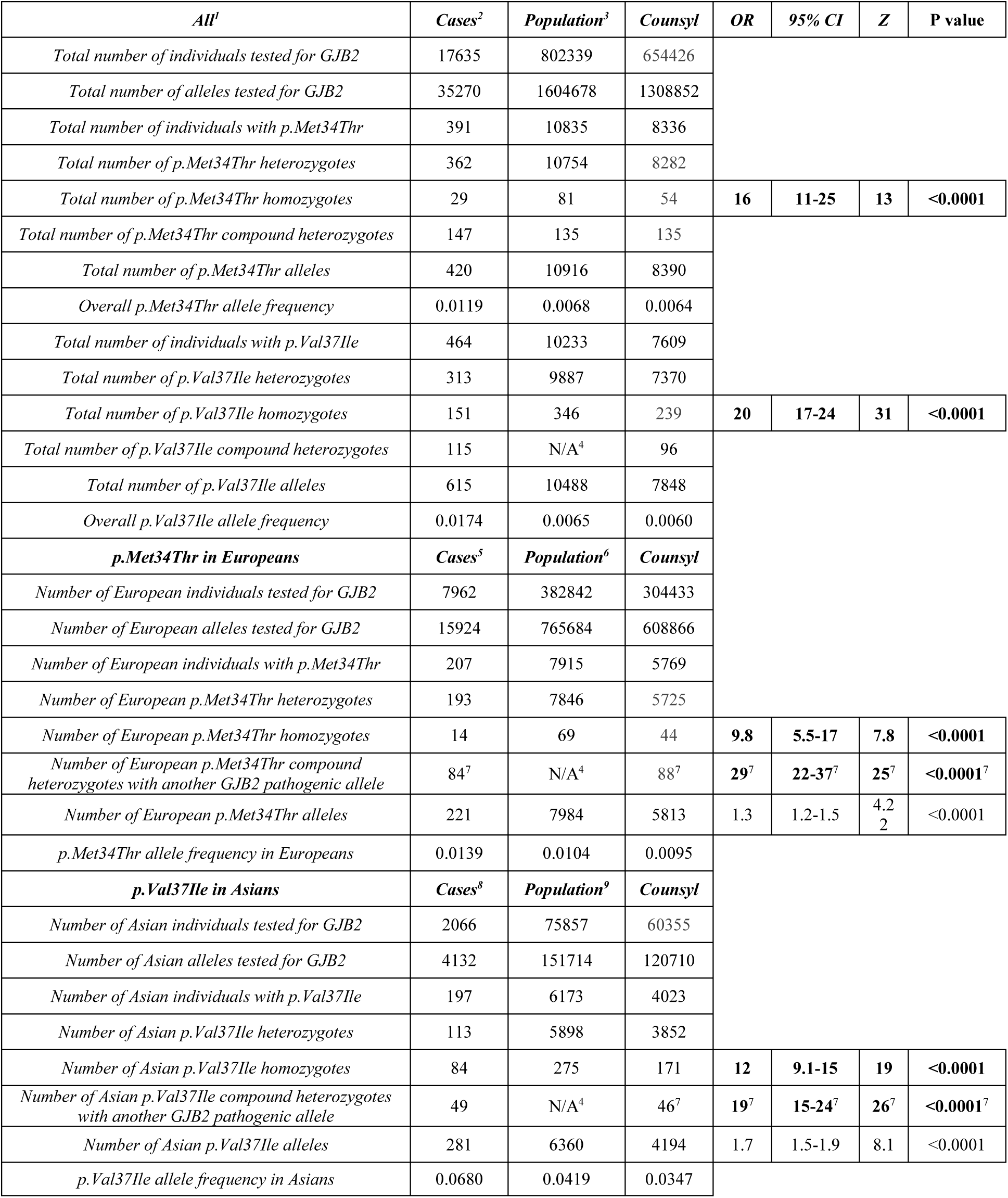
Summary statistics. 1. Only probands (unrelated individuals) were counted. However, we could not rule out the possibility of related cases from different sites because cases were de-identified before shared. Nevertheless, the likelihood of such occurrence would be low and would not significantly impact the conclusion.
2. The total number of cases included in statistical analyses does not include BCH, DY, and GDWC where the total number of individuals tested at these sites were not available.
3. The total population data were from Counsyl, CUHK, TAU, UNC, and gnomAD.
4. NA: Not available, because individual genotype information is not available from gnomAD.

Because the allele frequencies of Met34Thr and Val37Ile were observed highest in European and East Asian populations respectively, we performed statistical analysis on stratified subpopulations by ethnicity to remove the confounding factor of different ethnic compositions between cases and controls. The significance still holds when we performed ethnicity-specific analysis. Met34Thr homozygotes are significantly enriched in cases over the general population in Europeans (OR=9.8, 95%CI=5.5-17, Z=7.8, p<0.0001) and Val37Ile homozygotes Asians (OR=12, 95%CI=9.1-15, Z=19, p<0.0001), respectively (Table 2). When we consider all biallelic cases involving Met34Thr or Val37Ile, including homozygotes and compound heterozygotes with another pathogenic variant in *GJB2*, the enrichments are more significant with ORs of 29 (95%CI 22-37, Z=25, p<0.0001) for Met34Thr in Europeans and of 19 (95%CI 15-24, Z=26, p<0.0001) for Val37Ile in Asians in cases with hearing loss in our multicenter cohort over the general population who underwent carrier screening at Counsyl. Therefore, both variants meet PS4 (Prevalence in affected individuals statistically increased over controls) according to the ACMG/AMP guidelines^2^. In contrast, although the allele frequencies were statistically enriched in cases in our multicenter cohort over the general population with ORs of 1.3 (95%CI 1.2-1.5, Z=4.2, p<0.0001) for Met34Thr in Europeans and of 1.6 (95%CI 1.4-1.9, Z=8.1, p<0.0001) for Val37Ile in Asians, the ORs do not meet the criteria for PS4 (>5). We conclude that comparing genotype frequency is more appropriate than allele frequency when we apply PS4 to interpret variants associated with an autosomal recessive condition.

### Computational predictions

The REVEL scores are 0.702 and 0.657 for Met34Thr and Val37Ile, respectively. PP3 could be applied to Met34Thr (REVEL>=0.7) but neither PP3 nor BP4 could be applied to Val37Ile because its REVEL score does not reach the threshold (0.7) for PP3 but exceeded the ceiling (0.15) for BP4 as specified by the ClinGen HL-EP^14^.

Different amino acid changes at positions M34 and V37 have been reported in patients with hearing loss.

Available evidence suggested these variants are of uncertain significance but with some evidence to support pathogenicity (Supplementary Information). While they do not constitute sufficient evidence for PM5, multiple VUS towards pathogenicity affecting the same amino acid residues corroborate with each other. Therefore, we applied PM5_supporting to Met34Thr and Val37Ile according to the criteria recently specified by the ClinGen HL-EP^14^.

### Segregation Data

Literature review showed that both variants segregated with hearing loss in additional family members. Furthermore, data from contributing laboratories indicate at least 35 affected siblings with the same homozygous or compound heterozygous genotype, including 16 with Met34Thr and 21 with Val37Ile (Supplementary Table S2). Therefore, both variants meet PP1_Strong according to the ACMG/AMP guidelines^2^ and the criteria recently specified by the ClinGen HL-EP^14^.

It should be noted that Met34Thr and Val37Ile homozygotes are present in large population databases and identified by population carrier screening tests. However, clinical information was not available in these individuals. Because people with mild hearing loss may not be diagnosed, we cannot rule out the possibility that these individuals may be mildly affected. Furthermore, it has been reported that Val37Ile homozygotes lose hearing at approximately 1 dB per year^15^, suggesting an age-dependent penetrance of the hearing loss phenotype. The penetrance by young adulthood is estimated to be 17%^16^. Therefore, we do not consider children and adults of reproductive age with normal hearing and biallelic Met34Thr or Val37Ile as observations in controls (BS2) or non-segregation (BS4).

### Allelic Data

Both variants have been identified in affected homozygotes or compound heterozygotes. In our multicenter cohort, we observed 138 Met34Thr compound heterozygotes and 141 Val37Ile compound heterozygotes with other pathogenic/likely pathogenic alleles among 428 probands with hearing loss (Supplementary Table S2). However, because these variants are relatively common in the population and because we have sampled a large number cases, we only applied PM3 and did not upgrade the evidence to the strong level despite the large number of biallelic cases identified.

### Functional Data

The Met34Thr variant altered gap junction function in *Xenopus* oocytes^17-19^ and mammalian cells^20-22^. This variant had a similar expression pattern as wild-type connexin 26 in human sweat glands, in transfected HeLa cells, and in transfected COS-7 cells^18,20,21^. Paired *Xenopus* oocytes showed robust conductance when injected with *in vitro* transcribed human wild-type *GJB2* mRNA, reduced conductance when co-injected with wild-type and variant mRNA, and no coupling conductance above background when injected with variant mRNA only^17-19^. Co-injection of *GJB2* c.101T>C with wild-type connexin 30 (encoded by *GJB6*) or connexin 31 (encoded by *GJB3*) also showed reduced conductance^19^. Single channel conductance in c.101T>C transfected HeLa cells is reduced^22^, as supported by molecular dynamics simulations^23^. Furthermore, c.101T>C transfected HeLa cells did not transfer Lucifer yellow (a fluorescent dye) to neighbors across gap junctions compared to wild-type connexin 26 transfected cells^20,22,24^ and the ability to transfer neurobiotin was also reduced.^21^ However, c.101T>C transfected HeLa cells was able to load the dye in response to non-physiological zero extracellular Ca^2+^ stimulus^25^, suggesting that the Met34Thr variant connexin may retain some residual function under unusual circumstances. The atomic structure of the human connexin 26 gap junction channel revealed that the methyl group in Met34 in the first transmembrane domain interacts with the tryptophan at amino acid position 3 in the amino-terminal helix of an adjacent protomer to stabilize the hexameric channel^26^, and alteration of the residue is predicted to impact function^27^.

Similarly, Xenopus oocytes injected with mRNA encoding Val37Ile showed no conductance above background, and co-injection of Val37Ile variant connexin 26 with wild-type connexin 26, connexin 30, or connexin 31 showed reduced conductance^19,28^. Propidium iodide dye transfer was impaired in *GJB2* c.109G>A transfected HEK293T cells^29^. Homozygous knock-in mouse model of c.109G>A in *Gjb2* was reported to have progressive mild hearing loss, more pronounced at higher frequencies^30^. However, because the codon for Val37 in mouse (GTG) was different from that in human (GTT), the c.109G>A variant in mouse would translate into Val37Met. Nonetheless, this *in vivo* animal study indicates that alteration of Val37 impacts hearing.

These functional studies suggest Met34Thr and Val37Ile impact connexin 26 function; however, Xenopus oocytes and mammalian cell lines may not truly reflect the biologic process in human cochlea. Therefore, we applied PS3_Supporting to both variants according to the criteria recently specified by the ClinGen HL-EP^31^.

### Genotype-Phenotype Correlation

We obtained clinical information for 472 cases with biallelic *GJB2* genotypes involving Met34Thr or Val37Ile (Supplementary Table S2) including 183 with Met34Thr, 306 with Val37Ile, and 17 with [Met34Thr];[Val37Ile]. Met34Thr was *in trans* with a premature termination variant in 86 cases with clinical information, including 73 [Met34Thr];[Gly12fs] compound heterozygotes. Val37Ile was *in trans* with a premature termination variant in 92 cases with clinical information, including 32 [Val37Ile];[Leu79fs] compound heterozygotes. Although information was incomplete and non-uniform, we characterized hearing loss for different genotypes. Most cases had hearing loss before 18 years old. The hearing loss was typically bilateral, mild to moderate, and affecting mid- to high-frequencies. Progression of hearing loss was reported in all genotype categories (Table 3). In some individuals, profound hearing loss may be due to age-dependent progression or other etiologies. For example, one had meningitis at 1.5 years old, one was 76 years old when tested, and one infant had asymmetric hearing loss with moderate loss in the left ear and severe-to-profound loss in the right ear (Supplementary Table S2).

**Table 3.** Characteristics of hearing cases with biallelic *GJB2* variants involving p.Met34Thr or p.Val37Ilele, including homozygotes and compound heterozygotes with another pathogenic or likely pathogenic variant. Abbreviations: [P/LP], a pathogenic or likely pathogenic allele in *GJB2*; [PTC], an allele with a premature termination codon in *GJB2*; OR, odds ratio; CI, confidence interval; p, p value.

| Genotype | [Met34Thr];<br>[P/LP] | [Met34Thr];<br>[Met34Thr] | [Met34Thr];<br>[Val37Ile] | [Met34Thr];<br>[PTC] | [Val37Ile];<br>[P/LP] | [Val37Ile];<br>[Val37Ile] | [Val37Ile];<br>[PTC] | c.[35delG];<br>[35delG]<sup>1</sup> |
| --- | --- | --- | --- | --- | --- | --- | --- | --- |
| **Total probands** | 138 | 27 | 17 | 78 | 131 | 139 | 78 | 86 |
| **Total with characterization<sup>2</sup>** | 183 | 28 | 17 | 86 | 127 | 150 | 91 | 86 |
| **Age of Onset<sup>3</sup>** | 177 | 28 | 17 | 83 | 139 | 144 | 78 | 86 |
| Childhood (<18 years) | 155 (88%) | 23 (82%) | 16 (94%) | 72 (87%) | 113 (81%) | 120 (83%) | 50 (64%) | 84 (98%) |
| **Laterality** | 133 | 21 | 11 | 62 | 109 | 111 | 66 | 86 |
| Unilateral | 4 (3%) | 0 | 1 (9%) | 2 (3%) | 5 (5%) | 4 (4%) | 3 (5%) | 0 (0%) |
| Bilateral | 129 (97%) | 21 (100%) | 10 (99%) | 60 (97%) | 104 (95%) | 107 (96%) | 63 (95%) | 86 (100%) |
| Asymmetrical | 15 (12%) | 3 (27%) | 2 (20%) | 9 (15%) | 6 (6%) | 7 (7%) | 1 (2%) | 0 (0%) |
| **Frequency ranges<sup>4</sup>** | 66 | 14 | 6 | 30 | 59 | 59 | 34 | 1 |
| High | 41 (62%) | 9 (64%) | 5 (83%) | 13 (43%) | 35 (59%) | 41 (69%) | 21 (62%) | 1 (100%) |
| Mid | 17 (26%) | 4 (29%) | 1 (17%) | 11 (37%) | 3 (5%) | 10 (17%) | 2 (6%) | 0 (0%) |
| Low | 0 (0%) | 0 (0%) | 0 (0%) | 0 (0%) | 7 (12%) | 0 (0%) | 6 (18%) | 0 (0%) |
| **Progression** | 20 (11%) | 2 (7%) | 4 (24%) | 10 (12%) | 20 (16%) | 10 (7%) | 13 (14%) | 4 (4%) |
| **Severity** | 146 | 24 | 14 | 63 | 121 | 133 | 77 | 63 |
| Mild | 55 (38%) | 10 (42%) | 5 (36%) | 26 (41%) | 29 (24%) | 48 (36%) | 23 (30%) | 0 (0%) |
| Moderate | 67 (46%) | 8 (33%) | 7 (50%) | 29 (46%) | 55 (45%) | 52 (39%) | 34 (44%) | 7 (11%) |
| **Mild+Moderate** | **122 (84%)** | **18 (75%)** | **12 (86%)** | **55 (87%)** | **84 (69%)** | **90 (75%)** | **57 (74%)** | **7 (11%)** |
| Moderately severe | 12 (8%) | 3 (13%) | 2 (14%) | 3 (5%) | 20 (17%) | 12 (9%) | 9 (12%) | 9 (14%) |
| Severe | 4 (3%) | 0 (0%) | 0 (0%) | 3 (5%) | 16 (13%) | 20 (15%) | 10 (13%) | 9 (14%) |
| Profound | 8 (5%) | 3 (13%) | 0 (0%) | 2 (3%) | 1 (1%) | 1 (1%) | 1 (1%) | 38 (60%) |
| **Statistics (OR[95%CI], p)<sup>5</sup>** |  |  |  |  |  |  |  |  |
| [Met34Thr];[Val37Ile] |  | 0.86<br>[0.0-673],<br>0.97 |  |  |  |  |  |  |
| [Met34Thr];[PTC] |  | 0.76<br>[0.02-34],<br>0.89 | 0.88<br>[0.0-281],<br>0.97 |  |  |  |  |  |
| [Val37Ile];[Val37Ile] |  | 1.35<br>[0.04-49],<br>0.87 | 1.56<br>[0.0-432],<br>0.88 | 1.77<br>[0.47-7.7],<br>0.40 |  |  |  |  |
| [Val37Ile];[PTC] |  | 1.36<br>[0.04-57],<br>0.87 | 1.00<br>[0.34-3.00],<br>0.88 | 1.78<br>[0.33-9.4],<br>0.50 |  | 1.00<br>[0.34-3.00],<br>0.99 |  |  |
| c.[35delG];[35delG] |  | **79**<br>**[1.5-4210],**<br>**0.0315** | 91<br>[0.3-32548],<br>0.13 | **103**<br>**[11-902],**<br>**<0.0001** |  | **58**<br>**[10-336],**<br>**<0.0001** | **58**<br>**[7.7-438],**<br>**<0.0001** |  |

1. All information is not available for all individuals.
2. When age of onset was not available, age of testing was used as a surrogate.
3. Data from the Laboratory for Molecular Medicine only.
4. Flat audiogram shapes that affect all frequency ranges are under-reported.
5. Odds ratio refers to the genotype in the column header more likely to be milder than that in the row header. Statistical significant differences in severity are in bold.

Ordered logistic regression models showed no statistical significant difference in severity among Met34Thr or Val37Ile homozygotes, [Met34Thr];[Val37Ile], and their corresponding compound heterozygotes with a premature truncating variant (p=0.93), but Met34Thr or Val37Ile homozygotes or compound heterozygotes with a premature truncating variant presented significantly milder hearing loss than a cohort of 63 c.35delG homozygotes (p<0.05) from the Laboratory for Molecular Medicine, the majority of whom had profound hearing loss. [Met34Thr];[Val37Ile] compound heterozygotes also had milder hearing loss than c.35delG homozygotes, but the sample size was too small to reach statistical significance (Table 3). One infant with [Val37Ile];[c.235delC] passed newborn hearing screening, but was found to have unilateral mild high-frequency hearing loss at five weeks by audiology evaluation, and progressed to bilateral mild high frequency hearing loss at 20 weeks follow-up. One parent was self-reported to be unaffected, but after familial testing she was found to be [Met34Thr];[c.167delT], and audiology evaluation revealed mild high-frequency hearing loss at 47 years of age. This multicenter study identified five unaffected relatives with different genotypes via familial testing or population screening (Table 5). Interestingly, three were Asian and two were Ashkenazi Jewish. We also identified 411 individuals with biallelic Met34Thr (189) or Val37Ile (238) including 16 [Met34Thr];[Val37Ile] compound heterozygotes via carrier testing. However, their hearing status was not evaluated.

Taken together, phenotypic manifestation of *GJB2* Met34Thr and Val37Ile related hearing loss can vary from apparently normal to profound. However, these two variants are typically associated with bilateral mild to moderate sensorineural hearing loss, affecting high- to mid-frequencies. Progression has been reported. There is no significant difference in severity among Met34Thr or Val37Ile homozygotes and compound heterozygotes with another PTC variant (Table 3). The penetrance could not be calculated. However, among all individuals with biallelic *GJB2* variants involving Met34Thr or Val37Ile, more compound heterozygotes than homozygotes were observed in our multicenter case cohort compared to the carrier screening population by Counsyl for both Met34Thr (OR 2.03, 95%CI 1.22-3.37, Z=2.7, p<0.0064) and Val37Ile (OR 1.90, 95% 1.35-2.66, Z=3.7, p=0.0002), suggesting a higher penetrance for compound heterozygous genotypes than Met34Thr or Val37Ile homozygosity.

### Summary Interpretation

In summary, Met34Thr and Val37Ile can be classified as pathogenic based on PS4 (homozygotes and compound heterozygotes are significantly enriched in cases over the general population), PP1_Strong (segregated with hearing loss in many affected family members), PM3 (found *in trans* with many different pathogenic *GJB2* variants in patients with hearing loss), PM5_Supporting (multiple different amino acids changes at the same positions reported in patients with hearing loss), and PS3_Supporting (*in vitro* and *in vivo* evidence supporting functional impacts).

## DISCUSSION

### Interpreting variant using allele frequency information

The maximum population allele frequencies for Met34Thr and Val37Ile are 2% in Finnish and 8% in East Asian in gnomAD, respectively, and in 3.5% (7/198) of Finnish and 17% (32/186) of Chinese Dai in Xishuangbanna in 1000 Genomes Project, respectively. Because of the small sample sizes and because hearing status was not assessed in these reportedly healthy individuals, we cannot use these population frequencies as controls. The BA1 stand-alone rule (allele frequency is >5% in population databases) is not applicable here to Val37Ile, because there is conflicting evidence suggesting pathogenicity. Based on professional judgement, the ClinGen HL-EP ruled that PS4 overrides BA1.

When applying PS4, we need to choose appropriate cohorts of cases and controls. Because mild hearing loss may not be diagnosed without a formal audiology evaluation, we opted to compare frequencies in cases vs. the general population. Should the presence of the variants be significantly enriched in cases over the general population, they would be even more significantly enriched over normal controls.

In this study, we chose to compare the ratios of homozygotes and compound heterozygotes with Met34Thr or Val37Ile in cases vs. the general population. Because the majority of the alleles in the population were from heterozygous carriers who are not expected to have hearing loss due to their carrier status, the ORs of allele frequencies in our cases over the general population were significant (P<0.0001 and 95%CI of OR not including 1) but <5, thus not meeting PS4 according to ACMG/AMP guidelines^2^ (Table 2), consistent with a previous meta-analysis^32^. We observed a more significant effect when including compound heterozygotes, which could be explained by a higher penetrance of other pathogenic variants in *GJB2* than Met34Thr and Val37Ile. Our data indicate that it is preferable to include compound heterozygous individuals in the analysis. Because individual genotype data are not available from population databases such as gnomAD, population carrier screening data are invaluable.

We chose to use laboratory contributed data for cases instead of relying on published reports to avoid publication biases. Negative results are difficult to get published. Some patients may be included in multiple studies. Large cohort studies do not generally provide detailed case level data, and cases studies do not specify the total number of cases tested for the variant of interest.

Ethnicity information is crucial in case-control studies, because different composition of ethnicities could confound the results. In this study, we found the significances were inflated when ethnicity information was disregarded. The ratio of Met34Thr and Val37Ile homozygotes varies significantly from among laboratories due to different population compositions tested. Combining data from different laboratories while ignoring ethnicity information would bias towards the ethnic composition of the laboratory that contributed the most data. Stratifying patients by ethnicity significantly reduces the bias, but decreases the sample sizes. Combining data from laboratories around the world allowed us to perform unbiased analysis with sufficient statistic power. Unfortunately, ethnicity information is not always submitted to the testing laboratories, and some laboratories do not collect such information. Furthermore, the ethnicity in gnomAD was inferred based on the principal component analysis, which may not be the same as what patients self-reported. GnomAD tends to over-estimate major populations as people with a self-reported mixed ethnic background would likely be counted towards one of the major populations. Therefore, we recommend laboratories to collect ethnic information on the test requisition form and request patients and physicians to provide ethnic information when ordering genetic tests. This information would be extremely informative in conducting analyses like the one in this study.

### Genotype-phenotype correlation

Variable expressivity of Met34Thr and Val37Ile has been reported^16,33^. The hearing loss may be unilateral or bilateral, from mild to profound, affecting different frequency ranges, even in individuals with the same genotype. Val37Ile has also been associated with pathopoieia^34^ and sudden hearing loss^35^. There have been over 100 publications reporting individuals with these variants. However, case-level genotype and phenotype data were limited. This multicenter study report over 96% of biallelic Met34Thr or Val37Ile cases with bilateral hearing loss including a small percentage with asymmetric presentations, 84% with mild to moderate hearing loss, and a majority with high frequency loss, consistent with previous reports^36-39^. Individuals with profound hearing loss may have an alternate etiology such as infection and old age. Unilateral or asymmetrical hearing loss may progress to bilateral uniform.

The majority of diagnosed cases in our multicenter cohort had a pediatric onset of hearing loss. The findings could be due to ascertainment bias, because most individuals undergo genetic diagnosis are younger than 18 years old. Adults with mild hearing loss may not seek audiology evaluation or genetic testing. Progression of hearing loss was only reported in a small percentage of cases, because the progression was slow^15^ and long-term follow-up information was not available.

The penetrance of these two variants could not be assessed, because the hearing status was unknown in individuals with biallelic Met34Thr or Val37Ile in the general population. These individuals may be truly unaffected or have mild hearing loss that is undiagnosed. It has been estimated that the penetrance of Val37Ile homozygotes was 17% in children in China^16^, but the number could vary by age, ethnicity, and the allele *in trans* in compound heterozygotes. Our data suggest that the penetrance is higher in Met34Thr or Val37Ile compound heterozygotes than in corresponding homozygotes. Three Asian Val37Ile homozygotes and two Ashkenazi Jewish Met34Thr compound heterozygotes were confirmed unaffected by audiology evaluation with the oldest known age at testing of 30 years (Table 4).

**Table 4.**
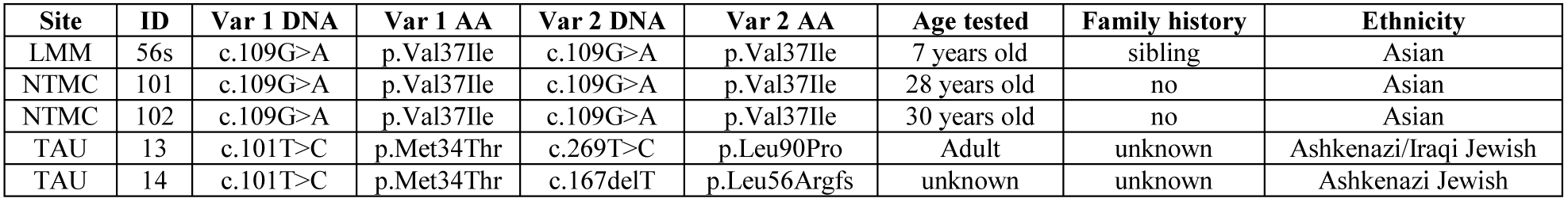
Unaffected Met34Thr or Val37Ile homozygotes or compound heterozygotes confirmed by audiology evaluation.

The mechanism for incomplete penetrance and variable expressivity remains elusive. Environmental and genetic modifiers may play a role. It is unclear how many different haplotypes were involved and whether the phenotype is haplotype-dependent. The haplotype with c.-684_-675del in 5’UTR of NM_004004.5 (rs139514105, referred to as c.-493del10) *in cis* with Met34Thr was confirmed in an affected sibpair from the UK^7^ and seven affected German individuals^40^, but its frequencies in all affected and unaffected biallelic Met34Thr populations are unknown. This haplotype did not abolish *GJB2* expression in cultured keratinocytes, which argues again a regulatory role of the noncoding cis variant^7^. Identification of the modifiers will not only help interpret the genetic findings but also point to effective therapeutic strategies for *GJB2*-related hearing loss, the most common form of hereditary hearing loss. Given the high allele frequency of these two variants in the population, further studies to identify the modifiers through population genetic screening and phenotypic evaluation of biallelic individuals are warranted and feasible.

### Barriers to accurate variant interpretation

Our experience in analyzing Met34Thr and Val37Ile in *GJB2* revealed barriers to accurate variant interpretation. First, no phenotypic information was available in large population studies or population screening. Although we were able to achieve statistical significance with a large sample size, should hearing status in the general population be known, a much smaller sample size would be sufficient to provide the same level of statistical power. Second, incomplete ethnicity, family history and clinical information from laboratories and publications diminished the usefulness of many cases. We would urge ordering physicians and testing laboratories to collect and share such information and editors and peer-reviewers to demand detailed case-level information for publication. Third, case information is biased towards published cases reports usually by clinicians and genetic service centers. Although they provide detailed clinical information, matched control information is mostly not available. A systematic patient registry to document harmonized clinical and genetic information will overcome this. Finally, given the slow progressiveness and variable expressivity, we still cannot accurately determine the penetrance of these variants in causing hearing loss and pinpoint the factors that influence the penetrance. Although age seems to be a factor, it could not explain everything. Long-term follow up of individuals with these variants in prospective studies will illuminate in this area.

## CONCLUSION

We demonstrate that the ACMG/AMP variant interpretation framework can be applied to variants with incomplete penetrance. Large and diverse sample sizes are required to overcome limitations and draw valid conclusions. The ClinGen HL-EP established a collaborative model of operation to collect case-level data, which allowed unbiased analysis of Met34Thr and Val37Ile, two controversial variants in *GJB2*. Based on compelling statistical and supporting functional evidence, we conclude that the two *GJB2* variants meet criteria to be classified as pathogenic for autosomal recessive sensorineural hearing loss, typically bilateral mild to moderate and slowly progressive over time^15^. When these variants are identified in homozygosity or compound heterozygosity with another pathogenic or likely pathogenic allele in *GJB2* in cases with severe to profound hearing loss, an alternate etiology should be investigated.

## ADDITIONAL INFORMATION

## Supporting information

## Author contributions

This work was overseen by the three co-chairs of the ClinGen Hearing Loss Working Group, SSA, HLR, and ANAT. AMO served as the coordinator of the working group. JS, AMO, IC, HD, TM, AP, HPK, SG, RM-H, KM, CL, MK, JA, YZ, YH, ML, YC, RC, Y-FC, KBA, GG, MAM-P, JG, YZ, KZ, ZB, LB-V, BD, MF, TW, NN, AW, SSA, and HLR contributed data to the study. JS, SEH, ARG, RKS, and MTD performed literature review. JS, XH, and ANAT performed data analysis. JS and ANAT drafted the manuscript. AMO, IC, HD, TM, HPK, RM-H, MK, YZ, YC, KBA, KZ, NN, and AW provided critical edits and feedback. Members of the ClinGen Hearing Loss Working Group participated in the discussion on applying ACMG/AMP and HL specific rules to determine the classification of the Met34Thr and Val37Ile variants. The final version was approved by all authors.

## Ethics statement

The study was approved by respective Institutional Review Boards, Helsinki Committees, or equivalent ethics committees of participating institutions involving research on human subjects.

---

Supplementary Information. Survey results, Met34Val and Val37Ile reported in various populations, analyses of missense variants at position Met34 and Val37, and Supplementary Table S1.

**Supplementary Table S2.** Clinical information of 472 hearing loss individuals with biallelic *GJB2* variants involving Met34Thr or Val37Ile.

## REFERENCES

1. Wang NK, Chiang JPW. Increasing evidence of combinatory variant effects calls for revised classification of low-penetrance alleles. Genetics in medicine: official journal of the American College of Medical Genetics. 2018.

2. Richards S, Aziz N, Bale S, et al. Standards and guidelines for the interpretation of sequence variants: a joint consensus recommendation of the American College of Medical Genetics and Genomics and the Association for Molecular Pathology. Genetics in medicine: official journal of the American College of Medical Genetics. 2015;17(5):405–424.

3. Kelsell DP, Dunlop J, Stevens HP, et al. Connexin 26 mutations in hereditary non-syndromic sensorineural deafness. Nature. 1997;387(6628):80–83.

4. Scott DA, Kraft ML, Carmi R, et al. Identification of mutations in the connexin 26 gene that cause autosomal recessive nonsyndromic hearing loss. Human mutation. 1998;11(5):387–394.

5. Scott DA, Kraft ML, Stone EM, Sheffield VC, Smith RJ. Connexin mutations and hearing loss. Nature. 1998;391(6662):32.

6. Wilcox SA, Saunders K, Osborn AH, et al. High frequency hearing loss correlated with mutations in the GJB2 gene. Human genetics. 2000;106(4):399–405.

7. Houseman MJ, Ellis LA, Pagnamenta A, et al. Genetic analysis of the connexin-26 M34T variant: identification of genotype M34T/M34T segregating with mild-moderate non-syndromic sensorineural hearing loss. Journal of medical genetics. 2001;38(1):20–25.

8. Cucci RA, Prasad S, Kelley PM, et al. The M34T allele variant of connexin 26. Genetic testing. 2000;4(4):335–344.

9. Kelley PM, Harris DJ, Comer BC, et al. Novel mutations in the connexin 26 gene (GJB2) that cause autosomal recessive (DFNB1) hearing loss. American journal of human genetics. 1998;62:792–799.

10. Abe S, Usami S, Shinkawa H, Kelley PM, Kimberling WJ. Prevalent connexin 26 gene (GJB2) mutations in Japanese. Journal of medical genetics. 2000;37(1):41–43.

11. Griffith AJ, Chowdhry AA, Kurima K, et al. Autosomal recessive nonsyndromic neurosensory deafness at DFNB1 not associated with the compound-heterozygous GJB2 (connexin 26) genotype M34T/167delT. American journal of human genetics. 2000;67(3):745–749.

12. Feldmann D, Denoyelle F, Loundon N, et al. Clinical evidence of the nonpathogenic nature of the M34T variant in the connexin 26 gene. European journal of human genetics: EJHG. 2004;12(4):279–284.

13. Pollak A, Skorka A, Mueller-Malesinska M, et al. M34T and V37I mutations in GJB2 associated hearing impairment: evidence for pathogenicity and reduced penetrance. American journal of medical genetics Part A. 2007;143A(21):2534–2543.

14. Oza AM, DiStefano MT, Hemphill SE, et al. Expert specification of the ACMG/AMP variant interpretation guidelines for genetic hearing loss. Human mutation. 2018;39(11):1593–1613.

15. Wu CC, Tsai CH, Hung CC, et al. Newborn genetic screening for hearing impairment: a population-based longitudinal study. Genetics in medicine: official journal of the American College of Medical Genetics. 2017;19(1):6–12.

16. Chai Y, Chen D, Sun L, et al. The homozygous p.V37I variant of GJB2 is associated with diverse hearing phenotypes. Clinical genetics. 2015;87(4):350–355.

17. White TW, Deans MR, Kelsell DP, Paul DL. Connexin mutations in deafness. Nature. 1998;394(6694):630–631.

18. Skerrett IM, Di WL, Kasperek EM, Kelsell DP, Nicholson BJ. Aberrant gating, but a normal expression pattern, underlies the recessive phenotype of the deafness mutant Connexin26M34T. FASEB J. 2004;18(7):860–862.

19. Palmada M, Schmalisch K, Bohmer C, et al. Loss of function mutations of the GJB2 gene detected in patients with DFNB1-associated hearing impairment. Neurobiol Dis. 2006;22(1):112–118.

20. Martin PE, Coleman SL, Casalotti SO, Forge A, Evans WH. Properties of connexin26 gap junctional proteins derived from mutations associated with non-syndromal heriditary deafness. Human molecular genetics. 1999;8(13):2369–2376.

21. Thonnissen E, Rabionet R, Arbones ML, Estivill X, Willecke K, Ott T. Human connexin26 (GJB2) deafness mutations affect the function of gap junction channels at different levels of protein expression. Human genetics. 2002;111(2):190–197.

22. Bicego M, Beltramello M, Melchionda S, et al. Pathogenetic role of the deafness-related M34T mutation of Cx26. Human molecular genetics. 2006;15(17):2569–2587.

23. Zonta F, Buratto D, Cassini C, Bortolozzi M, Mammano F. Molecular dynamics simulations highlight structural and functional alterations in deafness-related M34T mutation of connexin 26. Front Physiol. 2014;5:85.

24. D’Andrea P, Veronesi V, Bicego M, et al. Hearing loss: frequency and functional studies of the most common connexin26 alleles. Biochemical and biophysical research communications. 2002;296(3):685–691.

25. de Wolf E, van de Wiel J, Cook J, Dale N. Altered CO2 sensitivity of connexin26 mutant hemichannels in vitro. Physiol Rep. 2016;4(22).

26. Maeda S, Nakagawa S, Suga M, et al. Structure of the connexin 26 gap junction channel at 3.5 A resolution. Nature. 2009;458(7238):597–602.

27. Yilmaz A. Bioinformatic Analysis of GJB2 Gene Missense Mutations. Cell Biochem Biophys. 2015;71(3):1623–1642.

28. Bruzzone R, Veronesi V, Gomes D, et al. Loss-of-function and residual channel activity of connexin26 mutations associated with non-syndromic deafness. FEBS letters. 2003;533(1–3):79–88.

29. Kim J, Jung J, Lee MG, Choi JY, Lee KA. Non-syndromic hearing loss caused by the dominant cis mutation R75Q with the recessive mutation V37I of the GJB2 (Connexin 26) gene. Exp Mol Med. 2015;47:e169.

30. Chen Y, Hu L, Wang X, et al. Characterization of a knock-in mouse model of the homozygous p.V37I variant in Gjb2. Scientific reports. 2016;6:33279.

31. Oza A, DiStefano M, Hemphill S, et al. 2018.

32. Chan DK, Chang KW. GJB2-associated hearing loss: systematic review of worldwide prevalence, genotype, and auditory phenotype. The Laryngoscope. 2014;124(2):E34–53.

33. Lameiras AR, Goncalves AC, Santos R, et al. The controversial p.Met34Thr variant in GJB2 gene: Two siblings, one genotype, two phenotypes. International journal of pediatric otorhinolaryngology. 2015;79(8):1316–1319.

34. Dai ZY, Sun BC, Huang SS, et al. Correlation analysis of phenotype and genotype of GJB2 in patients with non-syndromic hearing loss in China. Gene. 2015;570(2):272–276.

35. Chen K, Sun L, Zong L, et al. GJB2 and mitochondrial 12S rRNA susceptibility mutations in sudden deafness. Eur Arch Otorhinolaryngol. 2016;273(6):1393–1398.

36. Lim LH, Bradshaw JK, Guo Y, et al. Genotypic and phenotypic correlations of DFNB1-related hearing impairment in the Midwestern United States. Archives of otolaryngology--head & neck surgery. 2003;129(8):836–840.

37. Lee KH, Larson DA, Shott G, et al. Audiologic and temporal bone imaging findings in patients with sensorineural hearing loss and GJB2 mutations. The Laryngoscope. 2009;119(3):554–558.

38. Kenna MA, Feldman HA, Neault MW, et al. Audiologic phenotype and progression in GJB2 (Connexin 26) hearing loss. Archives of otolaryngology--head & neck surgery. 2010;136(1):81–87.

39. Jiang Y, Huang S, Deng T, et al. Mutation Spectrum of Common Deafness-Causing Genes in Patients with Non-Syndromic Deafness in the Xiamen Area, China. PloS one. 2015;10(8):e0135088.

40. Zoll B, Petersen L, Lange K, et al. Evaluation of Cx26/GJB2 in German hearing impaired persons: mutation spectrum and detection of disequilibrium between M34T (c.101T>C) and -493del10. Human mutation. 2003;21(1):98.

