## Supplementary material for "Consensus interpretation of the Met34Thr and Val37Ile variants in *GJB2* by the ClinGen Hearing Loss Expert Panel"

**Supplementary Information**

**Survey results on interpretation of *GJB2* Met34Thr and Val37Ile**

The link of an online survey was sent to laboratories performing genetic testing on *GJB2* to provide classification of the Met34Thr and Val37Ile variants. Twelve clinical genetic testing laboratories from United States and Canada responded to the survey. The classifications of the variants were controversial. Results are shown in the bar graphs below.

c.101T>C p.Met34Thr

|  |  |
| --- | --- |
| Pathogenic | 4 |
| Pathogenic with reduced penetrance | 1 |
| Likely Pathogenic | 2 |
| Likely pathogenic with reduced penetrance and milder phenotype | 1 |
| VUS | 3 |
| VUS - Suspicious | 1 |

c.109G>A p.Val37Ile

|  |  |
| --- | --- |
| Pathogenic | 8 |
| Pathogenic with reduced penetrance and milder phenotype | 1 |
| Pathogenic with reduced penetrance | 1 |
| Likely pathogenic | 2 |

**Identification of Met34Thr and Val37Ile in various populations**

Met34Thr has been reported in Colombian^1^, Estonian^2^, Finnish^3,4^, French^5,6^, German^7^, Italian^8,9^, Polish^10^, Russian^11^, Yakut (of Sakha)^12^, Slovak^13^, and multiracial^14,15^ patients with hearing loss. Val37Ile has been reported in Chinese^16-19^, Yakut (of Sakha)^12^, Taiwanese^20^, Tibetan^21^, and multiracial^14,15^ patients with hearing loss.

**Analyses of *GJB2* missense variants at amino acid residues 34 and 37**

Both amino acid residues M34 and V37 are evolutionarily conserved in most vertebrates. However, each of the Met34Thr, Met34Leu, and Val37Leu changes are present in seven fish species. Most computation prediction tools (15/23 for Met34Thr and 16/23 for Val37Ile) suggest an impact to protein (Supplementary Table S1). However, Polyphen2 and CADD predict that the Met34Thr change is tolerated, while SIFT/PROVEAN and CADD suggest that Val37Ile is tolerable.

Met34Val was only reported in the heterozygous state in one Turkish individual with profound hearing loss^22^ but was absent from gnomAD. The REVEL score for this variant is 0.786, and is therefore a variant of uncertain significance (VUS) based on PM2 and PP3. M34L was reported in the heterozygous state in two Thai female siblings with profound hearing loss^23^ but was also identified in 1/17248 East Asian chromosomes by gnomAD. Its REVEL score is 0.611, and is classified as a VUS (PM2). Met34Arg was identified in the heterozygous state in one individual with hearing loss,^24^ but was absent from gnomAD, and with a REVEL score of 0.924. It is considered a VUS (PM2, PP3). Finally, Met34Ile was reported in compound heterozygosity with c.35delG in an individual with profound hearing loss^15^ and is absent in gnomAD. The REVEL score is 0.730 though this variant Met34Ile is still of uncertain significance (VUS) based on PM2, PM3, and PP3.

Val37Ala was identified in the heterozygous state in one Canadian aboriginal with moderate hearing loss who also had a heterozygous variant (NM_ 006783.4 (*GJB6*):c.631T>G/p.Cys211Gly) of uncertain significance in *GJB6*^24^, and in another individual without detailed information^25^, but a second pathogenic allele in *GJB2* was not identified in either case. It has been found in 13/34420 Latino and 8/24032 African chromosomes by gnomAD. Its REVEL score is 0.675, and is therefore a VUS (PM2_Supporting). Val37Leu was found in the heterozygous state in five individuals^24^, and its REVEL score is 0.723. It was not identified in gnomAD and is classified as a VUS (PM2).

Overall, the available evidence suggested other variants at Met34 and Val37 are of uncertain significance but with some evidence to support pathogenicity. While they do not constitute sufficient evidence for PM5, multiple VUS towards pathogenicity affecting the same amino acid residues corroborate with each other (Supplementary Table 1).

**References:**

1.	Tamayo ML, Olarte M, Gelvez N, et al. Molecular studies in the GJB2 gene (Cx26) among a deaf population from Bogota, Colombia: results of a screening program. International journal of pediatric otorhinolaryngology*.* 2009;73(1):97-101.

2.	Teek R, Kruustuk K, Zordania R, et al. Prevalence of c.35delG and p.M34T mutations in the GJB2 gene in Estonia. International journal of pediatric otorhinolaryngology*.* 2010;74(9):1007-1012.

3.	Lopponen T, Dietz A, Vaisanen ML, et al. Homozygous M34T mutation of the GJB2 gene associates with an autosomal recessive nonsyndromic sensorineural hearing impairment in Finnish families. Acta oto-laryngologica*.* 2012;132(8):862-873.

4.	Lopponen T, Vaisanen ML, Luotonen M, et al. Connexin 26 mutations and nonsyndromic hearing impairment in northern Finland. The Laryngoscope*.* 2003;113(10):1758-1763.

5.	Roux AF, Pallares-Ruiz N, Vielle A, et al. Molecular epidemiology of DFNB1 deafness in France. BMC Med Genet*.* 2004;5:5.

6.	Leclere JC, Le Gac MS, Le Marechal C, Ferec C, Marianowski R. GJB2 mutations: Genotypic and phenotypic correlation in a cohort of 690 hearing-impaired patients, toward a new mutation? International journal of pediatric otorhinolaryngology*.* 2017;102:80-85.

7.	Zoll B, Petersen L, Lange K, et al. Evaluation of Cx26/GJB2 in German hearing impaired persons: mutation spectrum and detection of disequilibrium between M34T (c.101T>C) and -493del10. Human mutation*.* 2003;21(1):98.

8.	Amorini M, Romeo P, Bruno R, et al. Prevalence of Deafness-Associated Connexin-26 (GJB2) and Connexin-30 (GJB6) Pathogenic Alleles in a Large Patient Cohort from Eastern Sicily. Ann Hum Genet*.* 2015;79(5):341-349.

9.	Cama E, Melchionda S, Palladino T, et al. Hearing loss features in GJB2 biallelic mutations and GJB2/GJB6 digenic inheritance in a large Italian cohort. Int J Audiol*.* 2009;48(1):12-17.

10.	Pollak A, Skorka A, Mueller-Malesinska M, et al. M34T and V37I mutations in GJB2 associated hearing impairment: evidence for pathogenicity and reduced penetrance. American journal of medical genetics Part A*.* 2007;143A(21):2534-2543.

11.	Barashkov NA, Pshennikova VG, Posukh OL, et al. Spectrum and Frequency of the GJB2 Gene Pathogenic Variants in a Large Cohort of Patients with Hearing Impairment Living in a Subarctic Region of Russia (the Sakha Republic). PloS one*.* 2016;11(5):e0156300.

12.	Barashkov NA, Dzhemileva LU, Fedorova SA, Maksimova NR, Khusnutdinova EK. [Connexin gene 26 (GJB2) mutations in patients with hereditary non-syndromic sensorineural loss of hearing in the Republic of Sakha (Yakutia)]. Vestn Otorinolaringol*.* 2008(5):23-28.

13.	Minarik G, Tretinarova D, Szemes T, Kadasi L. Prevalence of DFNB1 mutations in Slovak patients with non-syndromic hearing loss. International journal of pediatric otorhinolaryngology*.* 2012;76(3):400-403.

14.	Samanich J, Lowes C, Burk R, et al. Mutations in GJB2, GJB6, and mitochondrial DNA are rare in African American and Caribbean Hispanic individuals with hearing impairment. American journal of medical genetics Part A*.* 2007;143A(8):830-838.

15.	Snoeckx RL, Huygen PL, Feldmann D, et al. GJB2 mutations and degree of hearing loss: a multicenter study. American journal of human genetics*.* 2005;77(6):945-957.

16.	Liu Y, Ke X, Qi Y, Li W, Zhu P. Connexin26 gene ( GJB2): prevalence of mutations in the Chinese population. Journal of human genetics*.* 2002;47(12):688-690.

17.	Xiao ZA, Xie DH. GJB2 (Cx26) gene mutations in Chinese patients with congenital sensorineural deafness and a report of one novel mutation. Chin Med J (Engl)*.* 2004;117(12):1797-1801.

18.	Zhao FF, Ji YB, Wang DY, et al. Phenotype-genotype correlation in 295 Chinese deaf subjects with biallelic causative mutations in the GJB2 gene. Genetic testing and molecular biomarkers*.* 2011;15(9):619-625.

19.	Luo J, Bai X, Zhang F, et al. Prevalence of Mutations in Deafness-Causing Genes in Cochlear Implanted Patients with Profound Nonsyndromic Sensorineural Hearing Loss in Shandong Province, China. Ann Hum Genet*.* 2017;81(6):258-266.

20.	Wang YC, Kung CY, Su MC, et al. Mutations of Cx26 gene (GJB2) for prelingual deafness in Taiwan. European journal of human genetics : EJHG*.* 2002;10(8):495-498.

21.	Duan SH, Ma JL, Yang XL, Guo YF. Simultaneous multigene mutation screening using SNPscan in patients from ethnic minorities with nonsyndromic hearingimpairment in Northwest China. Molecular medicine reports*.* 2017;16(5):6722-6728.

22.	Yilmaz A, Menevse S, Bayazit Y, Karamert R, Ergin V, Menevse A. Two novel missense mutations in the connexin 26 gene in Turkish patients with nonsyndromic hearing loss. Biochem Genet*.* 2010;48(3-4):248-256.

23.	Kudo T, Ikeda K, Kure S, et al. Novel mutations in the connexin 26 gene (GJB2) responsible for childhood deafness in the Japanese population. Am J Med Genet*.* 2000;90(2):141-145.

24.	Putcha GV, Bejjani BA, Bleoo S, et al. A multicenter study of the frequency and distribution of GJB2 and GJB6 mutations in a large North American cohort. Genetics in medicine : official journal of the American College of Medical Genetics*.* 2007;9(7):413-426.

25.	Azaiez H, Chamberlin GP, Fischer SM, et al. GJB2: the spectrum of deafness-causing allele variants and their phenotype. Human mutation*.* 2004;24(4):305-311.

**Supplementary Tables.**

**Supplementary Table S1**. Computational predictions of p.Met34Thr, p.Val37Ile, and other missense variants affecting the same codons in *GJB2*.

The results were obtained from <http://varcards.biols.ac.cn/>. Met34Thr and Val37Ile are with highlighted backgrounds (deleterious predictions in yellow and neutral predictions in green). REVEL scores >=0.7 are in red bold.

Supplementary Table S1. Computational predictions of p.Met34Thr, p.Val37Ile, and other missense variants affecting the same codons in *GJB2.*

| **NM_004004.5 (*GJB2*):** | **Met34** | | | | | **Val37** | | |
| --- | --- | --- | --- | --- | --- | --- | --- | --- |
| ***c.*** | 100A>T | 100A>G | 101T>C | 101T>G | 102G>A | 109G>A | 109G>C | 110T>C |
| ***p.*** | Met34Leu | Met34Val | Met34Thr | Met34Arg | Met34Ile | Val37Ile | Val37Leu | Val37Ala |
| ***Deleterious_vs_all_algorithms*** | 14:23 | 16:23 | 15:23 | 19:23 | 18:23 | 16:22 | 17:23 | 18:23 |
| ***Damaging_score*** | 0.61 | 0.70 | 0.65 | 0.83 | 0.78 | 0.73 | 0.74 | 0.78 |
| ***SIFT_score*** | 1 | 0.001 | 0.027 | 0 | 0.002 | 0.717 | 0.743 | 0.203 |
| ***SIFT_pred*** | Tolerable | Damaging | Damaging | Damaging | Damaging | Tolerable | Tolerable | Tolerable |
| ***Polyphen2_HDIV_score*** | 0.006 | 0.664 | 0.038 | 0.985 | 0.681 | 1 | 1 | 1 |
| ***Polyphen2_HDIV_pred*** | Benign | Possibly_damaging | Benign | Probably_damaging | Possibly_damaging | Probably_damaging | Probably_damaging | Probably_damaging |
| ***Polyphen2_HVAR_score*** | 0.01 | 0.34 | 0.083 | 0.778 | 0.34 | 0.996 | 0.998 | 0.999 |
| ***Polyphen2_HVAR_pred*** | Benign | Benign | Benign | Possibly_damaging | Benign | Probably_damaging | Probably_damaging | Probably_damaging |
| ***LRT_score*** | 0 | 0 | 0 | 0 | 0 | 0 | 0 | 0 |
| ***LRT_pred*** | Deleterious | Deleterious | Deleterious | Deleterious | Deleterious | Deleterious | Deleterious | Deleterious |
| ***MutationTaster_score*** | 1 | 1 | 1 | 1 | 1 | 1 | 1 | 1 |
| ***MutationTaster_pred*** | Disease_causing | Disease_causing | Disease_causing | Disease_causing | Disease_causing | Disease_causing_automatic | Disease_causing | Disease_causing |
| ***MutationAssessor_score*** | 0.11 | 1.295 | 2.315 | 2.865 | 2.865 | 2.095 | 2.035 | 1.47 |
| ***MutationAssessor_pred*** | Neutral | Low | Medium | Medium | Medium | Medium | Medium | Low |
| ***FATHMM_score*** | -5.08 | -5.17 | -5.41 | -5.45 | -5.3 | -5.46 | -5.43 | -5.35 |
| ***FATHMM_pred*** | Damaging | Damaging | Damaging | Damaging | Damaging | Damaging | Damaging | Damaging |
| ***PROVEAN_score*** | -0.18 | -2.09 | -3.8 | -4.47 | -1.62 | -0.82 | -2.32 | -3.26 |
| ***PROVEAN_pred*** | Tolerable | Tolerable | Damaging | Damaging | Tolerable | Tolerable | Tolerable | Damaging |
| ***VEST3_score*** | 0.804 | 0.909 | 0.409 | 0.948 | 0.852 | 0.703 | 0.941 | 0.955 |
| ***VEST3_pred*** | Damaging | Damaging | Tolerable | Damaging | Damaging | Damaging | Damaging | Damaging |
| ***MetaSVM_score*** | 0.488 | 1.005 | 0.765 | 1.096 | 1.094 | 0.5 | 1.105 | 1.042 |
| ***MetaSVM_pred*** | Damaging | Damaging | Damaging | Damaging | Damaging | Damaging | Damaging | Damaging |
| ***MetaLR_score*** | 0.8 | 0.89 | 0.843 | 0.964 | 0.931 | 0.89 | 0.962 | 0.948 |
| ***MetaLR_pred*** | Damaging | Damaging | Damaging | Damaging | Damaging | Damaging | Damaging | Damaging |
| ***M_CAP_score*** | 0.62 | 0.48 | - | 0.528 | 0.613 | - | 0.21 | 0.389 |
| ***M_CAP_pred*** | Damaging | Damaging | - | Damaging | Damaging | - | Damaging | Damaging |
| ***CADD_score*** | 0.65 | 12.92 | 14.43 | 23.4 | 23.7 | 10.55 | 14.65 | 14.84 |
| ***CADD_pred*** | Tolerable | Tolerable | Tolerable | Damaging | Damaging | Tolerable | Tolerable | Tolerable |
| ***DANN_score*** | 0.767 | 0.986 | 0.97 | 0.98 | 0.995 | 0.867 | 0.943 | 0.974 |
| ***DANN_pred*** | Tolerable | Tolerable | Tolerable | Tolerable | Damaging | Tolerable | Tolerable | Tolerable |
| ***FATHMM_MKL_score*** | 0.934 | 0.932 | 0.923 | 0.946 | 0.884 | 0.96 | 0.97 | 0.978 |
| ***FATHMM_MKL_pred*** | Damaging | Damaging | Damaging | Damaging | Damaging | Damaging | Damaging | Damaging |
| ***Eigen_score*** | -0.286 | 0.166 | 0.06 | 0.485 | 0.25 | 0.342 | 0.236 | 0.195 |
| ***Eigen_pred*** | Tolerable | Damaging | Damaging | Damaging | Damaging | Damaging | Damaging | Damaging |
| ***GenoCanyon_score*** | 1 | 1 | 1 | 1 | 1 | 1 | 1 | 1 |
| ***GenoCanyon_pred*** | Damaging | Damaging | Damaging | Damaging | Damaging | Damaging | Damaging | Damaging |
| ***fitCons_score*** | 0.534 | 0.534 | 0.534 | 0.534 | 0.534 | 0.534 | 0.534 | 0.534 |
| ***fitCons_pred*** | Tolerable | Tolerable | Tolerable | Tolerable | Tolerable | Tolerable | Tolerable | Tolerable |
| ***GERP_score*** | 5.21 | 5.21 | 5.21 | 5.21 | 5.21 | 5.21 | 5.21 | 5.21 |
| ***GERP_pred*** | Conserved | Conserved | Conserved | Conserved | Conserved | Conserved | Conserved | Conserved |
| ***phyloP_score*** | 3.304 | 3.304 | 5.061 | 5.061 | 2.028 | 6.102 | 6.102 | 9.263 |
| ***phyloP_pred*** | Conserved | Conserved | Conserved | Conserved | Conserved | Conserved | Conserved | Conserved |
| ***phastCons_score*** | 1 | 1 | 1 | 1 | 1 | 1 | 1 | 1 |
| ***phastCons_pred*** | Conserved | Conserved | Conserved | Conserved | Conserved | Conserved | Conserved | Conserved |
| ***SiPhy_score*** | 15.379 | 15.379 | 15.379 | 15.379 | 19.121 | 19.121 | 19.121 | 15.379 |
| ***SiPhy_pred*** | Conserved | Conserved | Conserved | Conserved | Conserved | Conserved | Conserved | Conserved |
| ***REVEL_score*** | 0.611 | **0.786** | **0.702** | 0.924 | 0.73 | 0.656 | **0.723** | 0.675 |
| ***REVEL_pred*** | Damaging | Damaging | Damaging | Damaging | Damaging | Damaging | Damaging | Damaging |

**Supplementary Table S2**. Clinical information of 472 hearing loss individuals with biallelic *GJB2* variants involving Met34Thr or Val37Ile, including homozygotes or compound heterozygotes with a pathogenic or likely pathogenic allele.

1. Variants are described according to the Human Genome Variation Society recommendations (<http://varnomen.hgvs.org/)>. NM_004004.5 is the reference cDNA used.
2. Parental confirmation of trans configuration were not performed in most cases. Alleles 1 and 2 are assumed to occur *in trans*, because there is only one coding exon in *GJB2*. If a variant were to be in cis with Met34Thr or Val37Ile, it would be expected to co-segregate in most cases unless it has occurred *de novo* in recent generations, which would be rare.
3. A field is blank, if specific information is not available. A field is N/A, if the information is not available for all cases from a site.
4. Age groups: congenital hearing loss, birth; >birth to 1 year old, infant; >1 year to <18 years old, childhood; >=18 years old, adult.
5. Severity: <15 unaffected, 15-30 mild, 31-50 moderate, 51-70 moderately severe, 71-90 severe, >90 profound. Recorded severity is the worst of all frequency ranges of both ears.
6. Unilateral: one ear is unaffected; asymmetric: bilateral but two ears are different.
7. Frequency range affected: rising, low-frequency loss; U, mid-frequency loss; sloping, high-frequency loss. Many flat audiograms (affecting all frequencies) were not specified, thus under-reported.
